# Photocrosslinked mucoadhesive hyaluronic acid hydrogel for transmucosal drug delivery

**DOI:** 10.1101/2025.06.14.659729

**Authors:** Seth Asamoah, Martin Pravda, Eva Šnejdrová, Martin Čepa, Mrázek Jiří, Carmen Gruber-Traub, Vladimír Velebný

## Abstract

Drug delivery to the central nervous system (CNS) is primarily hindered by the blood-brain barrier (BBB). To address this, mucoadhesive formulations have been designed to prolong residence time at the application site. In this study, we comprehensively characterized the physicochemical and mucoadhesive properties of hyaluronic acid tyramine (HATA) photocrosslinked hydrogels using rheological methods, nanoindentation, contact angle goniometry, and advanced confocal microscopy. A novel parameter, photon count per pixel, was introduced through confocal microscopy to assess hydrogel stability and mucoadhesion on *ex vivo* porcine olfactory tissues. Crosslinked hydrogels (1% and 2% w/v) exhibited stable mucoadhesive properties, ranging between 16.5 and 18 photon counts per pixel, whereas uncrosslinked counterparts typical of classical nasal formulations showed significant photon count losses (71% and 50% for 1% and 2% HATA, respectively). Nanoindentation analysis revealed a correlation between photoirradiation time, effective Young’s modulus, and mucoadhesion, identifying one minute of irradiation as optimal across all concentrations tested. The optimized hydrogels demonstrated mucoadhesive forces of 0.263, 0.412, and 0.701 mN.mm^-2^, corresponding to Young’s modulus values of 1995, 2465, and 2985 Pa for 1%, 2%, and 3% w/v HATA, respectively. These results highlight the importance of crosslinking for enhancing hydrogel stability and mucoadhesion. Additionally, BSA-labelled rhodamine served as a model protein drug in low-swelling hydrogels for drug release studies, laying the foundation for further optimization in targeted nasal drug delivery systems.

**Graphical Abstract:** 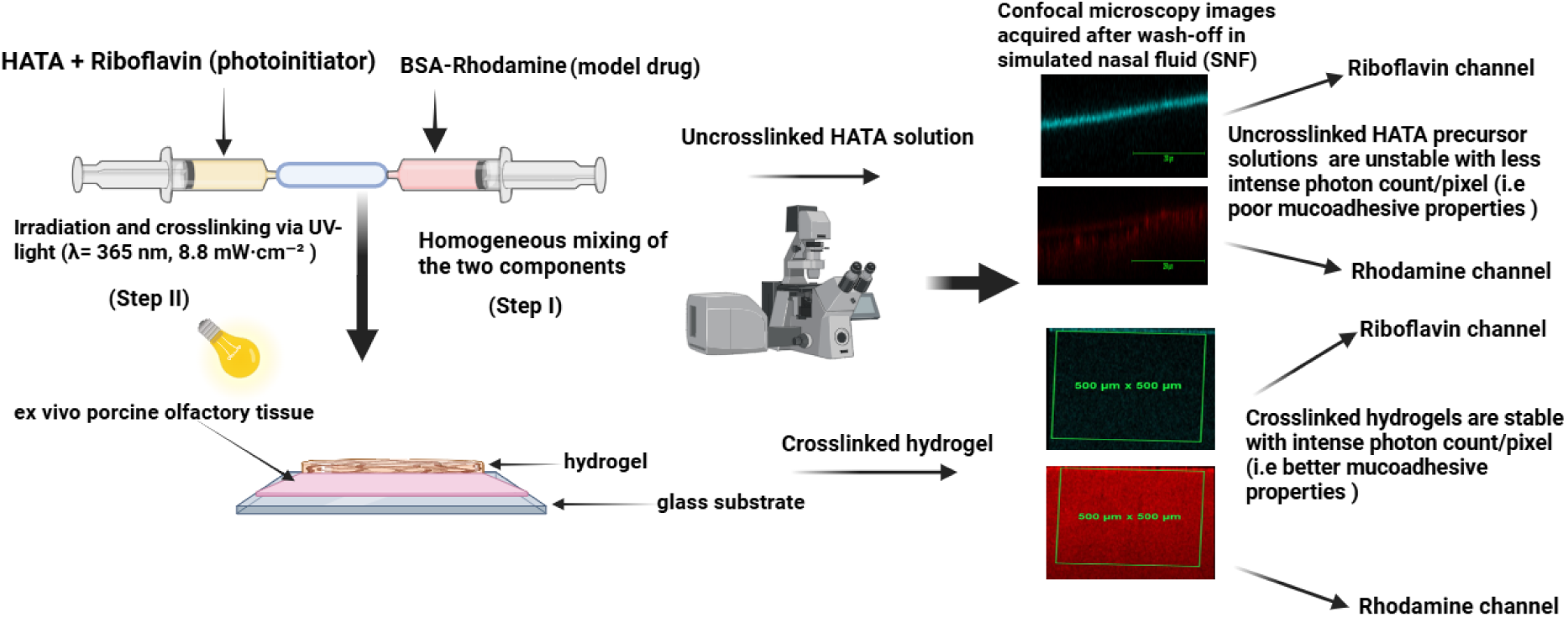

## 1 Introduction

In the treatment and management of central nervous system (CNS)-related diseases, a key challenge is achieving therapeutically effective drug concentrations due to the restrictive nature of the blood-brain barrier (BBB).[1] This barrier protects the brain from harmful substances while selectively allowing essential nutrients to pass through.[2] The intranasal route offers a non-invasive and efficient method for drug delivery to the brain by bypassing the BBB. This is made possible through its direct connection to the olfactory mucosa, located at the roof of the nasal cavity.[3] The nasal mucosa, being thin and porous, facilitates drug absorption, enabling both small molecules and biopharmaceuticals to reach the brain via the olfactory and trigeminal nerves. However, its thinness also leads to rapid drug clearance due to mucociliary activity, reducing the effectiveness of drug delivery. To address this, mucoadhesive formulations are employed to prolong drug residence time in the nasal cavity.[4,5]

The feasibility of the intranasal route of administration has been demonstrated in preclinical studies. For instance, intranasal administration of galantamine hydrobromide-loaded liposomes in rats showed enhanced acetylcholinesterase inhibition compared to oral administration.[6] This route is not limited to small-molecule drugs but also supports the delivery of peptides and proteins.[7] Given that several CNS-related disorders, such as Alzheimer’s and Parkinson’s disease, are linked to reduced insulin levels, intranasal insulin delivery has been explored in clinical trials. These studies have shown promising effects on learning, memory, energy regulation, and neuroprotection. Additionally, insulin-loaded gels formulated with carbopol and hydroxymethyl cellulose have been tested as a convenient alternative to injectable insulin.[8]

Despite these advantages, intranasal drug delivery faces challenges such as rapid mucociliary clearance and potential obstruction by highly viscous formulations.[9,10] To overcome these issues, *in situ* mucoadhesive drug delivery systems (MDDS) have been developed.[7]

Nasal MDDS are designed to deliver therapeutic agents to the mucosa-rich nasal cavity, ensuring effective local or systemic absorption. By adhering to the nasal mucosa, these systems prolong drug retention and enhance penetration, improving overall drug delivery efficiency.[11,12] By providing sustained drug release and avoiding first-pass metabolism, they offer a favourable environment for delivering proteins and peptides that are otherwise prone to gastrointestinal degradation. MDDS are formulated in various forms, including tablets, films, particles, and hydrogels.[13,14]

Hydrogels are 3-D systems usually made from different polymers with suitable crosslinking agent with the ability to imbibe and maintain large quantities of water.[15] Various polymers, such as cellulose derivatives, poloxamers [16], chitosan [17], tragacanth, carrageenan, xanthan gum [18] as well as hyaluronic acid (HA) [19] are commonly used in their formulation. HA, a linear glycosaminoglycan composed of N-acetyl-D-glucosamine and D-glucuronic acid, is particularly suitable due to its anionic surface charge, flexibility, and ability to form hydrogen bonds.[20,21] These properties contribute to its mucoadhesive and penetration-enhancing capabilities, making it a valuable component in drug delivery formulations.[22,23] HA has been explored for its mucoadhesive and penetration enhancement properties on mucosa surfaces based on its ability to form specific bonds with the mucin glycoproteins while prolonging the resident time of formulations on the surface. HA has been widely used in ophthalmology for dry eye treatment and in intranasal formulations, such as n-propyl gallate solid lipid nanoparticles, to improve mucoadhesive properties.[24] Intranasally, HA has been used for the formulation of n-propyl gallate solid lipid nanoparticles with improved mucoadhesive properties. [25] Additionally, its anti-inflammatory and protective effects have been beneficial in chronic rhinosinusitis treatments.[26]

For the scope of our work, *in situ* mucoadhesive UV-photocrosslinkable hydrogels based on a tyramine derivative of hyaluronic acid (HATA) have been developed. In a previous study, a nasal device with European patent number EP 3865173 A1 was developed for depositing substances onto the olfactory region through photocrosslinking, enabling the efficient delivery of therapeutics across the blood-brain barrier. This device provides a non-invasive and targeted method for delivering bioadhesive formulations to the nasal mucosa.[27] This study focuses on developing hydrophilic polymer formulations, particularly those based on hyaluronic acid (HA) and its derivative HATA, due to their strong mucoadhesive properties. Mucoadhesion is crucial for enhancing drug retention and interaction with the nasal mucosa, ensuring effective deposition. These properties suggest that the formulations could be well-suited for use with the nasal delivery device particularly the one mentioned here, improving drug delivery efficiency. Furthermore, HATA is particularly promising for drug delivery due to its superior and adjustable mechanical properties, which can be tailored to suit specific applications.[28,29] To initiate crosslinking, we used riboflavin, a vitamin B2 derivative known for its photoreactivity and low cytotoxicity, making it a clinically accepted and safe photoinitiator widely used in ophthalmic applications.[30] To the best of our knowledge, this is the first comprehensive physicochemical characterization of *in situ* photocrosslinkable mucoadhesive hydrogels based on HATA. We assessed both the precursor solution and the crosslinked hydrogel for stability and mucoadhesive properties using *ex vivo* porcine olfactory mucosal tissues. A novel confocal microscopy-based method was employed, measuring photon count per pixel to quantify hydrogel adhesion over time, providing direct evidence of mucoadhesion, unlike traditional wash-off methods. Additionally, a tack test was performed to correlate detachment force with the interfacial Young’s modulus, optimizing irradiation time while preserving mucoadhesive strength.

Ultimately, our photocrosslinked hyaluronic acid hydrogel formulation aims to enhance mucoadhesion, overcoming mucosal clearance challenges and improving nasal drug delivery efficiency.

## 2 Experimental section

### 2.1 Materials

Bovine serum albumin (BSA) was purchased from Sigma-Aldrich, Darmstadt, Germany. Porcine gastric mucin type II (PGM II), riboflavin, rhodamine B isothiocyanate-mixed isomers and low melting agarose were obtained from Sigma Aldrich, St. Louis, United States. HA (1 MDa) and its tyramine derivative HATA (1 MDa) with a degree of substitution 6.5% were supplied by Contipro a.s. Dolni Dobrouč, Czech Republic. Potassium dihydrogen phosphate was obtained from Lach-Ner s.r.o, Czech Republic. Sodium chloride, Sodium phosphate dibasic dodecahydrate and potassium chloride were purchased from Penta Chemicals Czech Republic.

### 2.2 Viscosity measurements

The viscosity of the PBS solutions of native HA and its derivative HATA at a concentration of 1, 2, and 3% w/v were evaluated on the DHR-3 rheometer (TA instruments, USA) using a 60 mm, 1° cone at a shear rate of 0.1-500 s^-1^ and a temperature of 25 □ C equipped with the TRIOS software. The flow curves obtained were fitted using the Carreau-Yasuda model.[31]

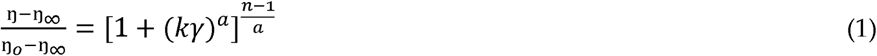

Where *η* is shear viscosity (Pa·s), *η*_∞_ is the infinite shear viscosity (Pa·s), *k* is the relaxation time (s), *n* is the power law index that characterizes the shear-thinning behavior, *η_0_* is the zero shear viscosity and *a*, is the term which describes the rate of transition from the Newtonian plateau to the power law region. The measurement was performed in triplicate.

### 2.3 Measurement of mucoadhesion by rheological synergism

For the evaluation of the mucoadhesive properties of the HA and HATA using the method of the rheological synergism [32], storage modulus *G’* (Pa) and loss modulus *G″* (Pa) of the solution of the polymers alone, the mucin alone, and the mixtures of mucin and polymers solutions were measured. For this purpose, 20% w/v dispersion of PGM II was freshly prepared by hydrating in PBS pH 6.5 and gently homogenized. 1 % and 2 % HA or HATA were dissolved in PBS pH 6.5 at a temperature of 60 □ C for 3 hours and left to stir overnight at room temperature. The equal volumes of polymer solution and PGM II dispersion were mixed and gently homogenized. The *G’* and *G″* were then evaluated on an absolute rotational rheometer KinexusPro+ (Malvern, United Kingdom) with a 40 mm upper plate geometry (PU 40) in oscillation mode via the amplitude sweep shear strain controlled with LVER from 0.1% and 100% shear strain at a frequency of 1 Hz. The measurement was conducted at 32 °C to mimic the physiological temperature of the nasal cavity.[33] The rheological synergism parameter was computed according to the equations (2) and (3). Measurements were performed in triplicates and averaged.

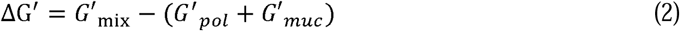

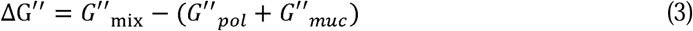

Where *G’_mix_*refers to the storage modulus of the mixture of mucin and HA or HATA, *G’_pol_*is storage modulus of polymer alone, *G’_muc_* is storage modulus of mucin alone, *G″_mix_* is the loss modulus of the mixture of mucin and HA or HATA, *G″_pol_* is storage modulus of polymer alone, *G″_muc_* is loss modulus of mucin alone.

### 2.4 Measurement of the wettability of the precursor solutions

1, 2 and 3% w/v of HATA or HA dissolved in PBS pH 7.4 were evaluated for their wettability on mucin-agarose coated glass. Mucin-agarose serving as a mucosal mimetic was produced according to the methods described by Spindler *et al* with modification. [34] Briefly, 3 % w/v low melting agarose was suspended in demineralized water at a temperature of 60 □ C under stirring conditions for 30 minutes to ensure complete dissolution. 10 % w/v of PGM II was also hydrated in demineralized water at 40 □ C until complete dissolution. The two components were then mixed in the ratio of 1:1 resulting in 1.5% agarose and 5% mucin dispersion. To produce the mucin-agarose coated substrate, the pristine glass slide was wiped with isopropyl alcohol to remove any debris or particulate matter. The glass was then dipped in the mucin-agarose dispersion, left to cool and solidify at 4 □ C, and utilized straight away.

The tested solutions dosed as 10 µL with the aid of syringe with an outer diameter of 1.83 mm were deposited on the mucin-agarose substrate. The contact angle was automatically measured on the Data Physics Instruments GmbH’s device (OCA 20, DataPhysics Instruments GmbH, Filderstadt, Germany) with inbuilt light source to guide the droplet and equipped with high resolution video camera to capture the spreading dynamics of the droplet in real time and evaluated by the OCA software. The contact angle was computed as the average contact angle of left and right.

### 2.5 Measurement of the kinetics of hydrogel formation

Precursor solutions of the tyramine derivative of HA (HATA) were prepared by dissolving three different concentrations 1%, 2%, and 3% w/v in PBS containing 50 µg/mL riboflavin as a photoinitiator. The polymer solutions were stirred continuously for 3 hours at 60□°C, followed by overnight stirring at room temperature. Crosslinking was performed in situ using a TA Instruments DHR-3 rheometer equipped with the OmniCure® S2000 Spot UV Curing System (Excelitas Technologies), which served as the ultraviolet (UV) light source. The UV light had an intensity of 8.8 mW·cm□^2^ and a wavelength of 365 nm, which corresponds to the excitation maximum of riboflavin.[35] Briefly, 500 µL of the precursor solution was placed on the lower geometry of the rheometer, using 20 mm diameter Peltier plates for both the upper and lower geometries. The kinetics of gelation was studied at a fixed displacement of 1.0 × 10□^3^ rad and an angular frequency of 6.28319 rad/s over a duration of 500 seconds. The gap between the plates was set to 1400 µm. All measurements were performed in triplicate at 25□°C.

### 2.6 Measurement of the viscoelastic properties of crosslinked hydrogels

The viscoelastic properties of the hydrogels were evaluated by subjecting circular-shaped samples (500 µL) to increasing strain and recording their mechanical response. The linear viscoelastic region (LVER) the range in which the material deforms elastically without structural failure was evaluated according to ISO 6721-10 and EN/DIN EN 14770 standards, using a 5% tolerance deviation criterion. To prepare the hydrogels, 500 µL of HATA precursor solutions at concentrations of 1%, 2%, and 3% w/v were transferred into 20 mm Teflon molds using a syringe and photocrosslinked with UV light at an intensity of 8.8 mW·cm□^2^ for 60 seconds. The LVER was then measured using an absolute rotational rheometer (KinexusPro+, Malvern, United Kingdom) equipped with 20 mm parallel plate geometry (PU 20) at a gap of 1400 µm. The test was performed at a constant frequency of 1 Hz, with strain applied in the displacement range of 0.003–4 rad, sampled at 10 points per decade. All measurements were conducted at 25□°C in accordance with ISO standard conditions (ISO 6721-10) for the rheological evaluation of hydrogels. Each experiment was performed in triplicate, and the results were averaged.

### 2.7 Measurement of Young’s modulus of hydrogels

Measurements of the Young’s modulus was performed on the Piuma nanoindenter with a spherical glass probe (Optics11Life). The 500 µL circle-shaped hydrogel was fixed to the bottom of a Petri dish with surgical glue. The sample was then immersed in a phosphate buffer saline of pH 7.4. The measurement started 10 minutes after immersion. Probe parameters: tip radius 266 µm, cantilever stiffness 0.56 N m-1. Indentation profile: “indentation control” mode; depth 4 µm, speed 5 µm s-1. The obtained indentation curves were processed in the manufacturer’s software Viewer (Hertz model; curve fit up to 80 % of the maximum load). A grid of 3 x 3 points with a step of 20 µm was measured and the results averaged.

### 2.8 Measurement of the mucoadhesive properties of hydrogels by tack test

For the measurement of the mucoadhesive properties of the crosslinked HATA hydrogels, 20% w/v porcine gastric mucin type II hydrated in PBS pH 6.5 was impregnated in an adhesive tape applied on both lower and upper PU 20 geometries of rheometer (KinexusPro+ Malvern, United Kingdom). The measurement was conducted at a temperature of 32 °C to mimic the physiological temperature of the nasal cavity. [36] A modified axial test for the determination of tack and adhesion included in the rSpace for Kinexus software was used.[37] A circle-shaped hydrogel sample with a diameter of 20 mm was placed in the centre of the lower geometry. The gaping speed was set to 10 mm· s^-1^, the contact force was 0.25 N, and the contact time was 20 s. Adhesion was expressed as the maximum detachment force per contact area (mN·mm^-2^) and the measurements were performed in quadruplicate and averaged.

### 2.9 Conjugation of rhodamine B isothiocyanate to HATA

Rhodamine B isothiocyanate was conjugated to photocrosslinkable HATA via the reaction of the hydroxyl groups of the HATA with the isothiocyanate of the rhodamine according to methods previously reported with modification. [38] Briefly, 1g of HATA was dissolved in NaHCO_3_; pH 9 adjusted using NaOH. The polymer was allowed to solubilize for 3 hours at 60 ° C and left to stir overnight at room temperature. 50 mg of Rhodamine B isothiocyanate was added and left stirring at room temperature for 24 hours while been kept in a dark place. The mixture was then dialyzed in demineralized water for several days until all unreacted rhodamine was removed. The product was then freeze-dried, and the amount of conjugated dye was estimated by UV-VIS spectrophotometry. Rhodamine content of the HATA was estimated by UV-VIS spectroscopy as 9 µg/ml of polymer and used in the confocal microscopy studies.

### 2.10 Conjugation of rhodamine B isothiocyanate to BSA

Rhodamine B isothiocyanate was conjugated to BSA according to methods already described elsewhere with modification. [38] Briefly, 500 mg of BSA was dissolved in NaHCO_3_ at pH 9 adjusted using NaOH. After complete solubilization of the BSA, 25 mg of Rhodamine B isothiocyanate was pre-dissolved in 1:3 dimethylformamide and NaHCO_3_ it was then added and left stirring at room temperature for 24 hours while been kept in a dark place. The resulting solution was then dialyzed against demineralized water for several days until all unreacted dye was removed. The resulting product was then freeze-dried and used as a model protein for the *in vitro* release experiments.

### 2.11 Measurement of mucoadhesion by confocal microscopy

To measure the mucoadhesion of hydrogels or uncrosslinked polymer solutions, tissue was obtained following the methods already described elsewhere. [39] Briefly 200 µL precursor solution of HATA-Rhodamine (1% or 2% w/v) was gently immobilized on porcine olfactory mucosa tissue sourced from a local abattoir pigs. This was then irradiated with UV light of intensity 8.8 mW·cm^-2^ for 1 minute. Uncrosslinked polymer solutions were transferred onto porcine olfactory mucosa tissue and used as control representative of classical nasal formulations. The hydrogels or the uncrosslinked precursor solutions were then taken through several washing procedures by simulated nasal fluid (SNF) prepared by the dissolution of sodium chloride (2.1925 g), calcium chloride (0.145 g) and potassium chloride (0.745 g) into 250 ml of distilled water and the pH adjusted to 6.5 [40]. The washing volume of SNF was 10 ml. Fluorescence images were acquired after every washing step using Leica TCS SP8 X confocal microscope (Leica Microsystems) equipped with a Leica HC PL APO CS objective (10x/0.40 dry). The excitation and emission wavelengths were 405 nm and 420–520 nm for riboflavin, and 540 nm, and 550 – 595 nm for rhodamine. Images were acquired with 2 µm steps in photon counting mode for further quantification and in z-stacks covering the whole thickness of the sample. Simultaneously the reflection image was collected using 670 nm laser line. For quantification, we used Las X software to create orthogonal sections from the z-stacks and measured the mean fluorescence intensity. If the fluorescence layer was washed out, we instead used the maximum projection image of the z-stack to get the mean fluorescence intensity. The number of photons per pixel as a new parameter to describe both stability and mucoadhesive properties of formulations was introduced.

### 2.12 Measurement of the swelling properties of hydrogels

The swelling properties of HATA hydrogels were evaluated by fully immersing crosslinked circle-shaped samples with a diameter of 20 mm in 9.5 ml of PBS with pH 7.4 for 96 h. At specified time intervals, the hydrogels were removed, and excess water dried using tissue paper and noted the weight. The swelling ratio (%) was evaluated according to equation 4.

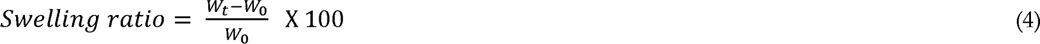

where *W_0_*is the hydrogel’s weight prior to immersion and *W_t_* is the hydrogel’s weight at specified time interval.

### 2.13 Model protein release study

The 1 % w/v HATA hydrogel was selected to test the drug release properties due to its fulfilment of various criteria for good mucoadhesion drug delivery system based on it spreadability and wettability, suitable viscosity and both *in vitro* and *ex vivo* mucoadhesive properties previously measured. The hydrogels were crosslinked as described previously. Briefly precursor solution of the 1% w/v HATA containing 250 µg/ml of BSA-Rhodamine conjugate as model protein was crosslinked for 1 minute and left overnight for complete hydrogel formation. The hydrogel was fully immersed in a 2 ml (PBS pH 7.4) as the release media and placed in an incubator (Witeg Wisd Incubator, WITEG Labortechnik) at 37 °C and a relative humidity of 96 %. At set time intervals all the medium was aspirated and replaced with a new one. The amount of BSA-Rhodamine released was determined to be quantified at 557 nm on the UV-VIS spectrophotometer. Calibration curve based on BSA-Rhodamine was established in the range of 3.9 µg/ml to 500 µg/ml with equation of a line of y = 0.0028 x – 0.0059, R^2^ = 0.9999. The release curve was fitted using the mathematical model developed by Korsmeyer-Peppas.

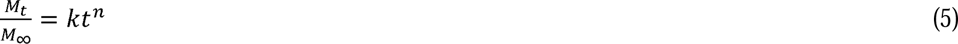

Where 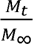 refers to the fraction of the drug released at time *t, k* is the release rate constant and *n* is the release exponent representative of the mechanism of drug release.

## 3 Results and discussion

Drug delivery systems (DDS) based on the tyramine derivative of hyaluronic acid (HATA) provide photocrosslinkable hydrogels when exposed to UV light. This method is advantageous due to the absence of toxic chemical reagents and leads to DDS with tuneable mechanical properties. Additionally, HATA hydrogels create a relatively conducive physiological environment for cells and other biomolecules to thrive. [41] In our work, we have developed HATA hydrogels to serve as carriers for the incorporation of APIs intended for nose to brain delivery.

### 3.1 Viscosity of the HA and HATA solutions

Viscosity of the formulation plays an indispensable role in intranasal DDS. In the formulation of *in situ* nasal gels, the precursor solution should meet prerequisites such as its ability to establish contact with the nasal mucosa and prevent dripping prior to crosslinking. The precursor solution should also have a viscosity low enough to promote its spreading and the overall mucoadhesive properties. [34] For the evaluation of viscosity, three different concentrations namely 1, 2, and 3% w/v of HATA or HA were chosen to check the suitability of a particular concentration of polymer for the development of the formulation. All the precursor solutions evaluated exhibited non-Newtonian flow behaviour as shown in Figure 1. with decrease in viscosity as the shear rate increased. The decrease in viscosity with increasing shear rate can be associated with the disruption of intramolecular hydrogen bonds, coupled with hydrophobic effects due to the reorganization and subsequent breakdown of polymer chains, eventually leading to a reduced viscosity of the polymer solutions. [42–44] As expected, there was a concentration dependent increase in viscosity with native HA exhibiting higher viscosities compared to the tyramine modified HA (HATA) an observation which we have associated with the chemical modification of the polymer. [45]

**Figure 1.**
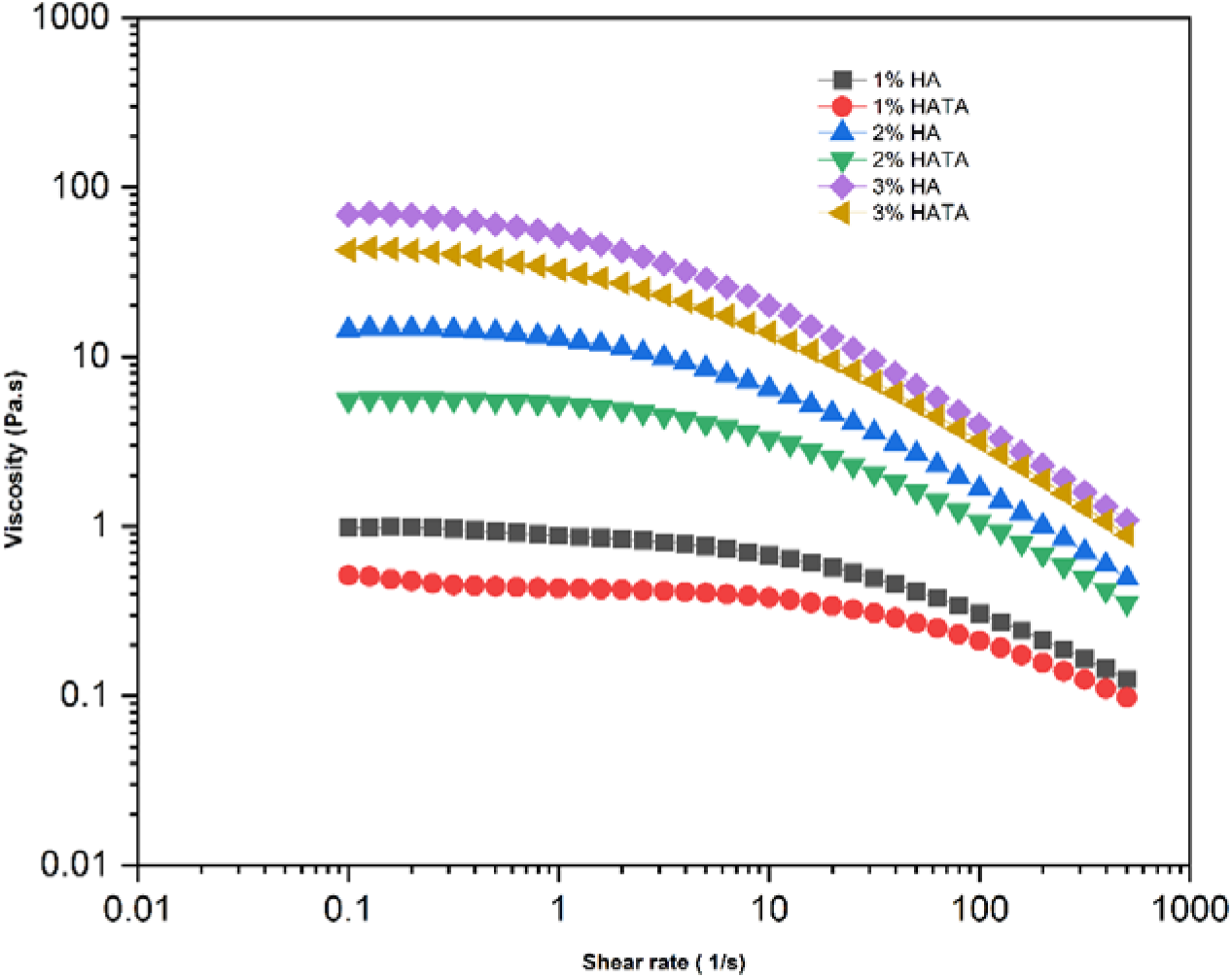
Viscosity of native hyaluronic acid (HA) and tyramine derivative of hyaluronic acid (HATA)

To fully understand the flow properties of these precursor solutions, we subjected the rheological data to the Carreau-Yasuda model previously described in equation (1). The model predicted a concentration dependent increase in the zero-shear viscosity (Table 1) of the solutions with native HA having higher zero shear rate viscosities compared to HATA despite their comparable molecular weight as detected by size exclusion chromatography. HA in its native form exist as highly organized secondary and tertiary structures which is perturbed during the chemical modification eventually affecting the polymer’s behaviour in solution. [46] This perturbation might have led to the drop in viscosity of the chemically modified form (HATA).

**Table 1.** Carreau-Yasuda model prediction of the zero-shear rate viscosity. The reported results is the average value of three measurements (Mean ± SD, n = 3).

| Polymer concentration | HA (Pa. s) | HATA (Pa. s) |
| --- | --- | --- |
| 1 % w/v | 1.03 ± 0.01 | 0.48 ± 0.04 |
| 2 % w/v | 15.56 ± 0.10 | 5.77 ± 0.01 |
| 3 % w/v | 77.58 ± 0.03 | 49.57 ± 2.21 |

### 3.2 Mucoadhesive properties of the precursor solutions determined by rheological synergism

Several theories have been proposed to describe the complex nature of mucoadhesion. Among them, the most common is the interdiffusion/interpenetration theory. This theory capitalizes on the formation of physical entanglements between polymer chains, mucin glycoproteins and secondary chemical interactions [47,48]. Rheology was used to evaluate the parameter of the rheological synergism. Generally, mucin-polymer mixture (muc+pol_mix) exhibited positive rheological synergism for both native HA in Figure 2A and the tyramine modified HATA in Figure 2B compared to the theoretical sum (muc+pol) of the individual components for the storage and loss moduli respectively. In both rheological parameters, HA exhibited the highest rheological values compared to the HATA due to highly ordered secondary and tertiary structures formed by the polymer as already depicted by the high viscosity observed in the native HA (Table 1). Despite the prevalence of a positive rheological synergism, there was an overall dominance of the viscous (*G″*) nature of the mixture over the elastic (*G’*) nature of the mixture (Figure 2A and 2B). This phenomenon can be explained by the inability of the mucin-polymer system to form a crosslinked gel. When the polymer concentration was raised from 1% to 2% w/v there was an increase in both *G’* and *G″* values as shown in Figure 2.

**Figure 2.**
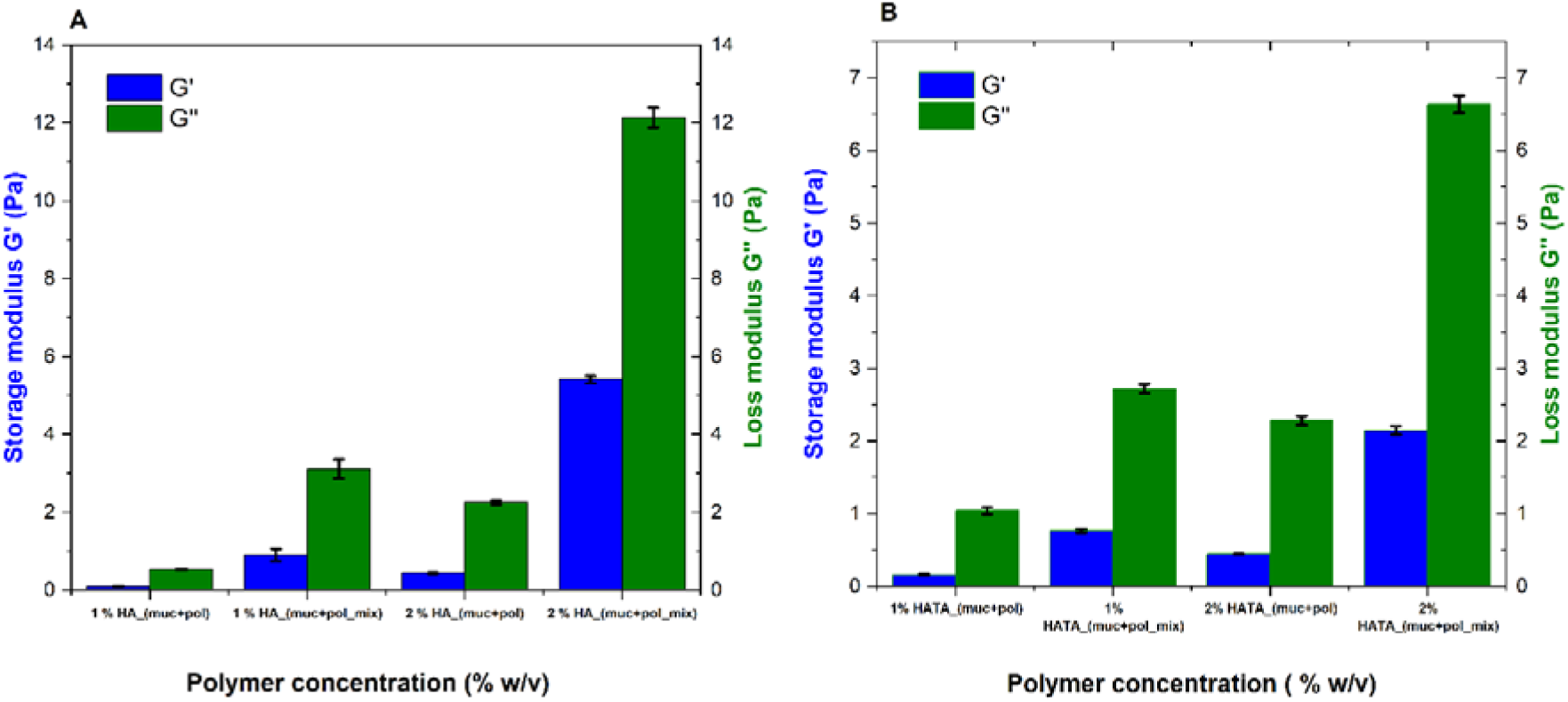
Evaluation of rheological synergism (A) HA: storage modulus (G’) and loss modulus (G″) (B) HATA: storage modulus (G’) and loss modulus (G″). Mucin-polymer mixture is designated as (muc+pol_mix) and theoretical sum of mucin and polymer is designated as (muc+pol)

Despite the two polymers having similar molecular weights, native HA exhibited a higher overall synergistic effect compared to its chemically modified counterpart, HATA. This difference is likely due to the structural alterations introduced during the modification process, which may affect HATA’s ability to interact with mucin. While native HA showed favorable properties in the synergism experiments, the crosslinkable nature of HATA offers significant advantages for drug delivery applications. Its tunable gelation properties and enhanced retention upon UV crosslinking demonstrated by photon-count confocal microscopy results in Section 3.7 make HATA a more efficient candidate for in situ nasal drug delivery systems.

### 3.3 Wettability of the precursor solutions

The wetting theory of mucoadhesion is demonstrated through the measurement of the contact angle. As the first step in the mucoadhesion process, it emphasizes the capacity of the bioadhesive polymer to spread and establish intimate contact with its substrate. According to the theory, the closer the contact angle approaches zero, the better the mucoadhesive properties. [22] To evaluate the wettability and spreadability of the precursor solutions and to select a suitable concentration, three different polymer concentrations, namely 1%, 2%, and 3% w/v of HA or HATA, were screened for their ability to establish contact and spread easily on the mucosa surface prior to crosslinking.

HA and HATA of similar molecular weight (1 MDa) were compared at the same concentrations to elucidate their spreading dynamics. Generally, the spreading dynamics of the precursor solutions displayed a concentration dependence, whether chemically modified or not (Figure 3). However, the native form (HA) exhibited relatively higher contact angles, most likely due to its highly ordered secondary and tertiary structure, which holds the polymer chains together, preventing easy spreadability. This observation agrees with the viscosities measured in Table 1, where higher viscosity values were recorded for the native form of the polymer. Upon the initial droplet deposition, designated as t0, from the needle, neither 1% nor 3% w/v of the native HA or the modified form (HATA) of the polymer produced significantly different contact angles (Figure 3A and 3C). However, HATA produced relatively lower contact angles with respect to the 2% w/v concentration of the polymer. The modified form of the polymer generally spreads more easily, with a stronger affinity to the hydrophilic mucosa model used, a feature required to achieve optimal mucoadhesive formulations. This easy spreadability and wettability of the polymers on the substrate further highlights the importance of using hydrophilic polymers, such as hyaluronic acid and its derivative HATA, in the development of nasal and mucoadhesive formulations.

**Figure 3.**
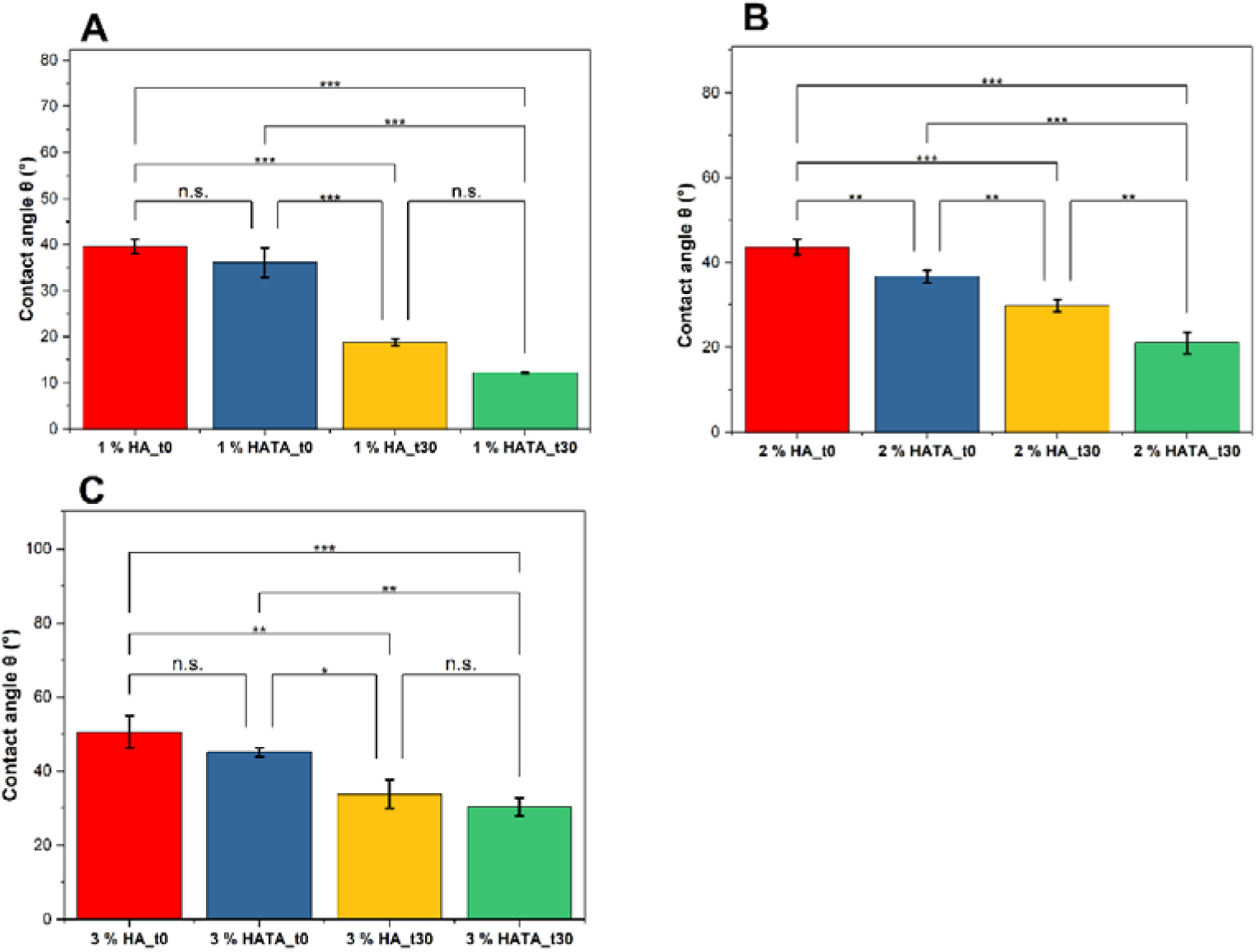
Contact angle measurement on mucin-agarose substrate as a function of time and concentration of native HA and modified HATA. (A). 1 % w/v (HA_t0, HA_t30) and 1 % w/v (HATA_t0, HATA_t30) (B). 2 % w/v (HA_t0, HA_t30) and 2 % w/v (HATA_t0, HATA_t30) (C). 3 % w/v (HA_t0, HA_t30) and 3 % w/v (HATA_t0, HATA_t30). t0 and t30 refers to contact angles measured at 0 and 30 seconds respectively after droplet deposition on the mucin agarose mucosal model.

### 3.4 Kinetics of hydrogel formation

The process of gelation of hydrogel precursor solutions was studied in real time to estimate the exact time where sol-gel transformation occurs under the irradiation of UV light at an intensity of 8.8 mW·cm^-2^. During this process, the photoinitiator riboflavin is transformed from its short-lived singlet state to the long-lived triplet state which then triggers the formation of di-tyramine from the tyramine derivative of HA. [49] For the evaluation of the kinetics of gelation, 1, 2, and 3% w/v of HATA precursor solutions were evaluated. In Figure 4, representative of a typical rheological diagram, the gelation time and crossover modulus, indicating the time of hydrogel formation and its strength respectively are extracted.

**Figure 4.**
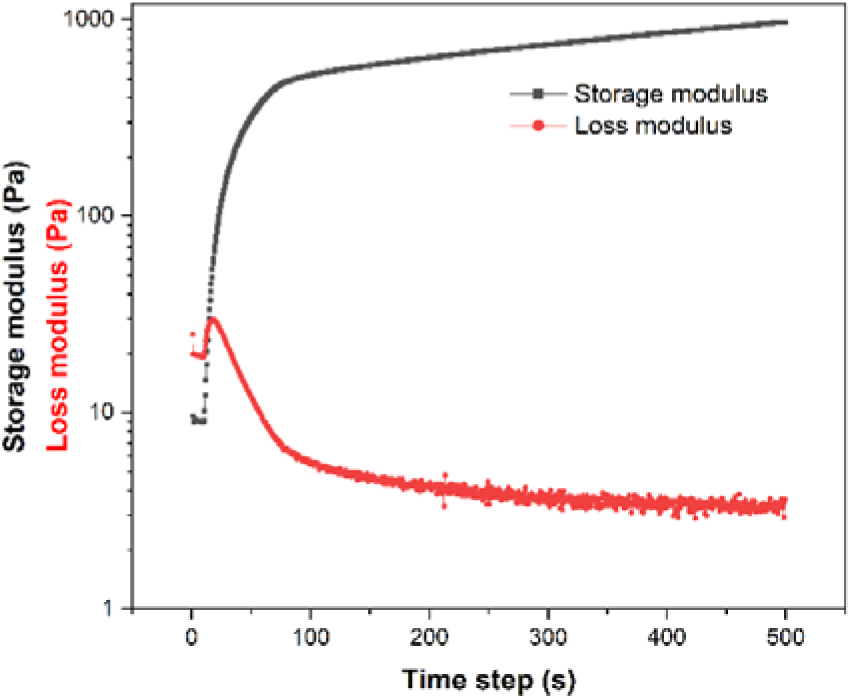
Typical rheological diagram of kinetics of gelation of HATA precursor solution.

All the three different concentrations of precursor solutions produced hydrogels at a very fast rate between 3 to 4 seconds as shown in Table 2. However, with increasing HATA concentration, there was a corresponding increase in the strength of the hydrogels designated as the crossover modulus due to the increase in the number of photocrosslinkable tyramine groups.

**Table 2.**
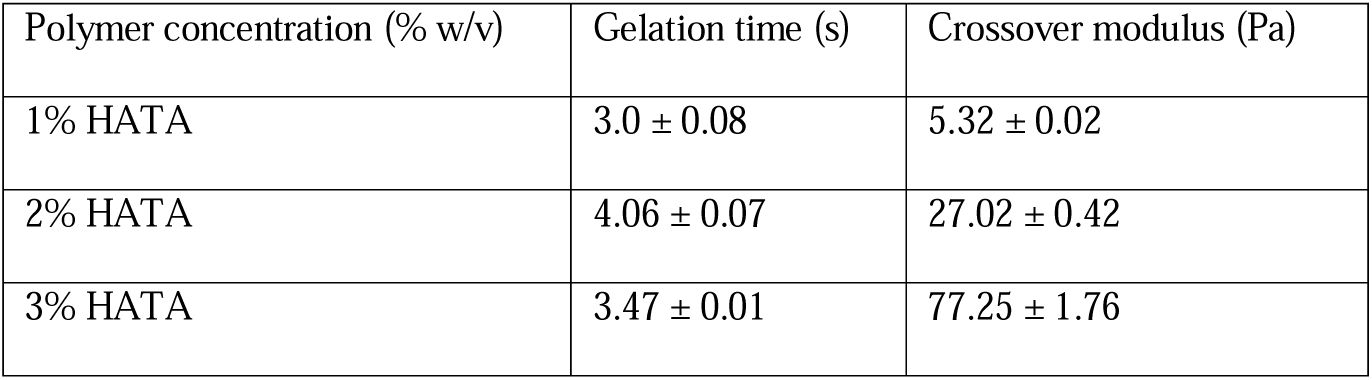
Kinetics of gelation of HATA at different concentrations and their corresponding crossover modulus. The reported results are the average value of three measurements (Mean ± SD, n = 3).

### 3.5 Effect of irradiation time on hydrogels properties

To determine optimal conditions for producing photocrosslinked mucoadhesive hydrogels, we investigated how varying UV irradiation time affects both the mechanical and mucoadhesive properties of the hydrogels. HATA precursor solutions at 1%, 2%, and 3% w/v were irradiated for 20, 60, or 180 seconds with UV light at an intensity of 8.8 mW·cm□^2^. The resulting hydrogels were then evaluated for mucoadhesion via tack testing and mechanical strength via nanoindentation (Figure 5).

**Figure 5.**
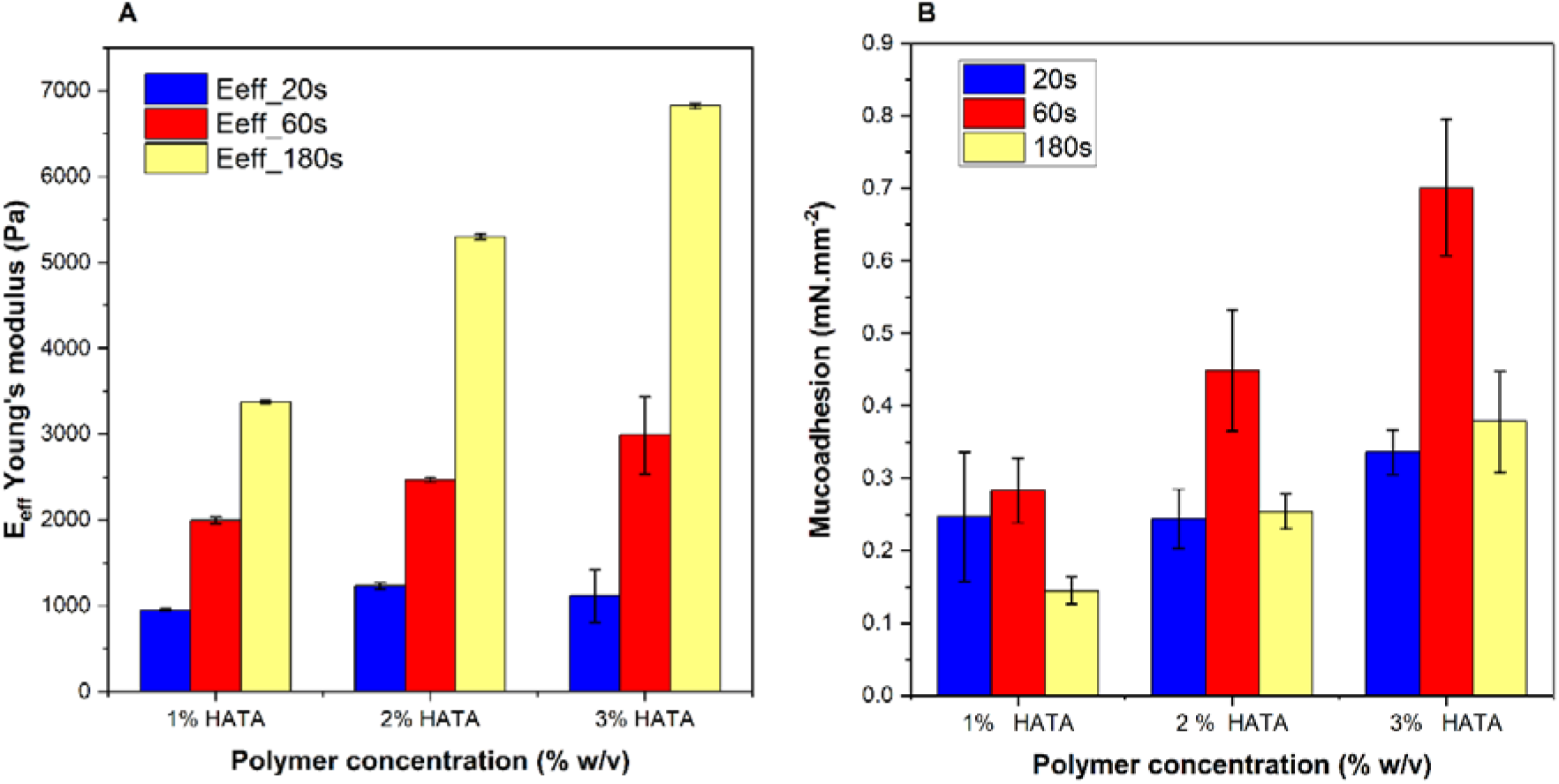
A) Irradiation time as a function of hydrogel’s Effective Young’s modulus (Eeff). 20s, 60s and 180s represent irradiation time in seconds B) Irradiation time as a function of mucoadhesion. 20s, 60s and 180s represent irradiation times.

The tack test, a standard method to assess adhesive behavior, measures the peak normal force required to detach the hydrogel from a mucosal substrate. This force was interpreted as the hydrogel’s mucoadhesiveness (Figure 5B). Simultaneously, we assessed the effective Young’s modulus (ELff) using a nanoindentation-based technique that provides non-destructive characterization of the hydrogel’s surface mechanical properties (Figure 5A). Unlike classical rheometry, this approach allows correlation between surface stiffness and mucoadhesiveness. [50,51]

Hydrogel strength, as indicated by ELff, increased with longer irradiation across all concentrations. At each time point, 3% HATA hydrogels showed the highest stiffness, followed by 2% and 1%. This trend emphasizes the need to balance crosslinking duration and polymer concentration to avoid over-crosslinking, which can reduce mucoadhesion by making the hydrogel too rigid (Figure 5A & 5B).

The relationship between mechanical strength and adhesion was further supported by the Dahlquist criterion, which states that materials with a storage modulus below 10□ Pa are effective adhesives due to their ability to deform and establish intimate contact with mucosal surfaces. [52] All tested hydrogels met this criterion, particularly at 60 seconds of irradiation, where sufficient stiffness was achieved without compromising mucoadhesiveness. As controls, uncrosslinked HATA_SOL formulations (1– 3% w/v) were also evaluated. These samples, which form physical hydrogels upon hydration, exhibited the highest tackiness due to the free movement and entanglement of polymer chains with mucin glycoproteins. The results, presented in Table 3, show that tackiness increased with concentration, confirming the influence of polymer content on adhesion in the uncrosslinked state. However, they suffer from key drawbacks such as low mechanical strength, susceptibility to enzymatic degradation [53], and instability at the application site, as confirmed by our newly developed confocal microscopy imaging technique (Figure 7).

**Figure 7.**
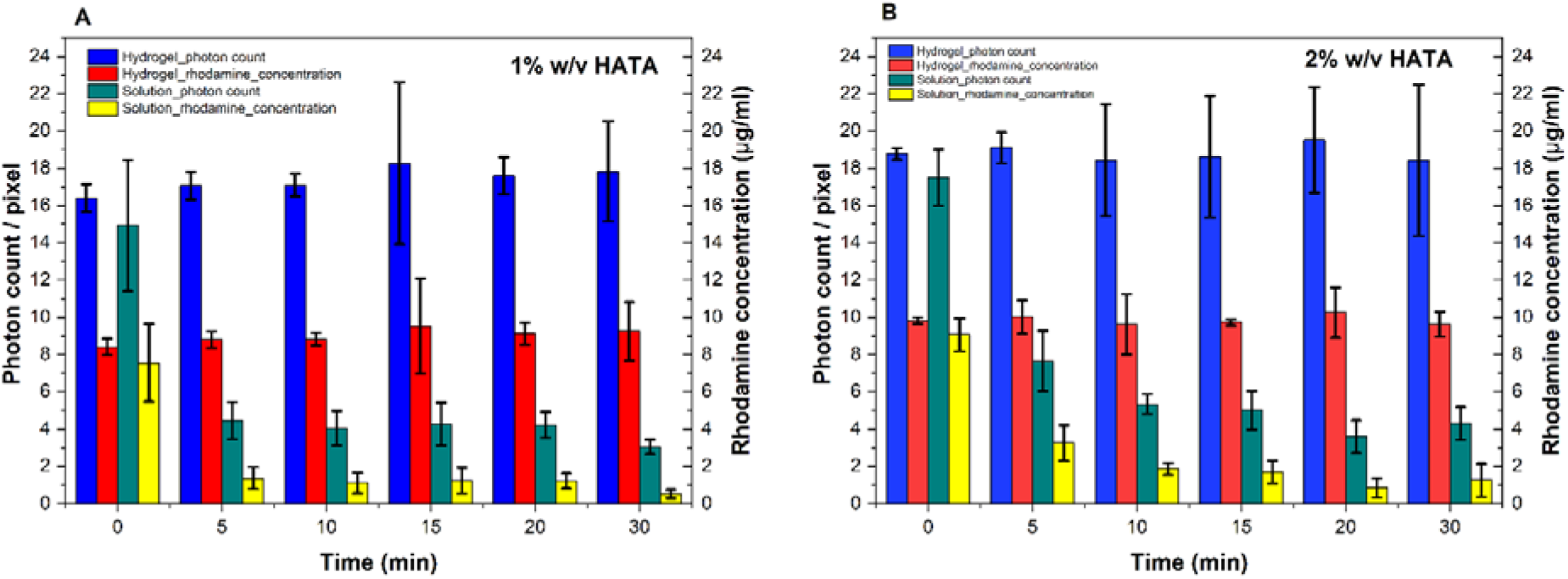
Photon count per pixel and corresponding rhodamine concentration for hydrogels and uncrosslinked solutions as a function of wash-off time in 1% w/v HATA (A) and 2% w/v HATA (B)

**Table 3.** Mucoadhesion (peak normal force) of crosslinked HATA hydrogels at three different concentrations (1%, 2%, and 3% w/v) irradiated for the optimized time of 60 seconds, compared to uncrosslinked HATA_SOL formulations used as controls. Results are reported as mean ± standard deviation (n = 4).

|  |  |
| --- | --- |
| Polymer concentration (% w/v) | Mucoadhesion (mN.mm<sup>-2</sup>) |
| 1% HATA | $0.263 \pm 0.029$ |
| 1% HATA_SOL | $2.55 \pm 0.45$ |
| 2% HATA | $0.412 \pm 0.049$ |
| 2% HATA_SOL | $5.85 \pm 1.225$ |
| 3% HATA | $0.701 \pm 0.094$ |
| 3% HATA_SOL | $7.022 \pm 1.758$ |

Overall, the crosslinked hydrogels displayed concentration-dependent mucoadhesive properties, with 3% HATA exhibiting the highest adhesion, followed by 2% and 1% (p < 0.05). These findings are consistent with those reported by Graça et al. [37], who demonstrated increased mucoadhesion with higher polymer concentrations in HA-based ocular delivery systems.

### 3.6 Viscoelastic properties of hydrogels

Precursor solutions of 1, 2 and 3 % w/v of HATA were irradiated for 60 seconds and their viscoelastic properties of the bulk hydrogel were determined using the rotational rheometer. The storage (G’) and loss moduli (G’’) depicting the elastic and viscous nature respectively of the hydrogels, were obtained from the rheological diagram in Figure 6 (see supplementary materials Table S1 for the absolute values). All three concentrations examined depicted a dominance of storage modulus signifying a complete formation of hydrogels. The hydrogel’s strength increased as the concentration of the precursor solution increased from 1 to 3% w/v. This phenomenon can be associated with the increase in crosslinked di-tyramine moieties as the concentration of the polymers increases.[54]

**Figure 6.**
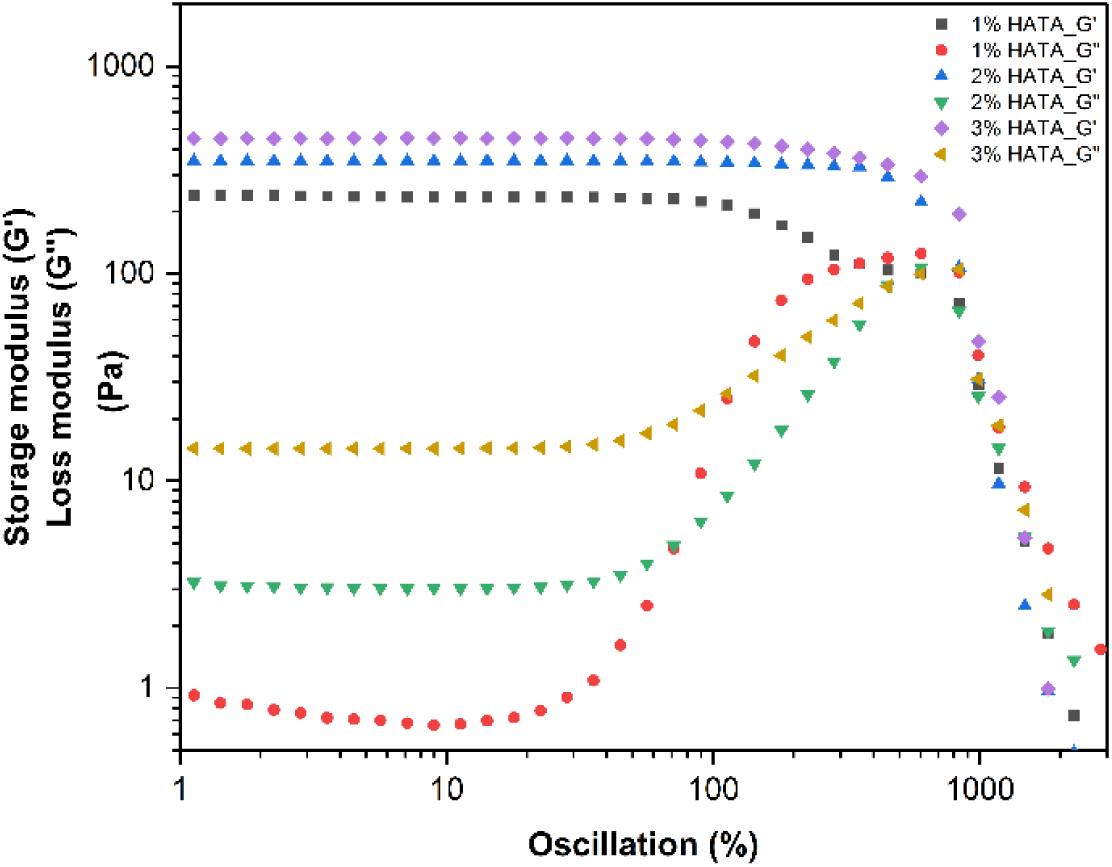
Linear viscoelastic region (LVER) of the three different concentrations of HATA after 60 seconds of irradiation

### 3.7 Evaluation of hydrogel stability and mucoadhesion by confocal microscopy

Imaging techniques such as atomic force microscopy, florescence microscopy and confocal laser scanning microscopy have been used in the past to characterize formulations for its mucoadhesive properties. [55] Imaging techniques are advantageous in their ability to provide direct quantification of adhered material on the mucosa surface. [55] For hydrogels, it qualifies as a suitable method in its ability to provide extra information such as the stability of the formulation at the site of application.

Hydrogel stability and adhesion on *ex-vivo* porcine olfactory tissue were evaluated by confocal microscopy. To highlight the importance of crosslinking the formulation at the site of application, the behaviour of both hydrogels and the uncrosslinked precursor solutions were evaluated. We introduced the novel parameter designated as the photon count per pixel to quantify the intensity of the covalently conjugated rhodamine-HATA hydrogel or solution on the *ex vivo* porcine olfactory mucosa tissue. We observed that crosslinked hydrogels were stable and were retained on the mucosa surface during entire thirty minutes of the wash-off experiments despite the complete removal of the riboflavin photoinitiator. We observed that the image intensity (see supplementary material Figure 1S) of the riboflavin decreased throughout the wash-off process while the covalently conjugated rhodamine-HATA signal remained relatively similar (see supplementary material Figure 1S). No statistically significant differences (see supplementary material Figure 2S) in stability and mucoadhesive properties were found between concentrations of 1% and 2% w/v rhodamine-HATA (Figure 7). The hydrogel photon count per pixel and corresponding rhodamine concentration remained constant throughout the washing process hovering between 16.5 to 18 photon count per pixel and 8 to 10 µg/ml respectively for both concentrations of crosslinked hydrogels tested (Figure7). This observation further confirms the successful conjugation of the rhodamine on the HATA. On the other hand, for the uncrosslinked solution of the polymer, 1% HATA demonstrated a drastic loss of about 71 % of its photon count, equivalent to the material loss or instability on the mucosa surface as demonstrated in Figure 7 and a thin layer of uncrosslinked material on the mucosal surface (see supplementary material Figure 1SB). This loss in material resulted in a corresponding decrease in the rhodamine concentration from about 8 µg/ml to 0.5 µg/ml at the end of the washing process.

For the 2% w/v HATA (Figure7B) of uncrosslinked material, about 50 % of the material was lost after 5 minutes of the washing process in comparison to the 1% w/v HATA (Figure 7A) in which 70 % of material was lost. This is because of the higher polymer concentration which eventually leads to higher viscosity and adhesion delaying the rapid removal of the material from the mucosa surface. This observation further highlights the importance of crosslinking the precursor solution to form a stable hydrogel rather than simply administering the formulation in a relatively low viscous uncrosslinked form.

### 3.8 Comparison of Mucoadhesion Characterization Methods

The mucoadhesion properties of the hydrogels were extensively characterized using various physicochemical methods, including contact angle goniometry, confocal microscopy, and tack test coupled with nanoindentation. Each method provides complementary insights into different aspects of mucoadhesion, offering a robust evaluation of the complex nature of the concept of mucoadhesion.

Contact angle goniometry was employed to assess the wettability and spreadability of the hydrogels on the mucin-agarose substrate, key indicators of their ability to establish intimate contact with the mucosal surface. A lower contact angle signifies better wettability, which is essential for strong mucoadhesive interactions, as it facilitates the polymer chains interpenetration with mucosal glycoproteins. This method allowed us to correlate polymer concentration and structural modifications with the spreading behavior of the precursor solutions, highlighting the improved wettability of the chemically modified hydrogels (HATA) compared to their native counterpart (HA). Confocal microscopy provided a novel approach to directly quantify the adhered hydrogels using photon count per pixel, offering a clear advantage over classical wash-off methods, which rely on indirect quantification methods. This method enabled us to visualize and confirm the stability of the hydrogels on *ex vivo* mucosal tissues.

In addition, tack test coupled with nanoindentation was used to evaluate the tackiness and the local interfacial viscoelastic properties (Effective Young’s modulus) of the hydrogels. Both techniques allowed us to establish a correlation between polymer concentration, irradiation duration, and the resulting mechanical properties, allowing optimization of hydrogel formulations without compromising mucoadhesion. By combining these methods, we obtained a comprehensive understanding of mucoadhesion from both mechanical and physicochemical perspectives. However, each method has its limitations. For instance, results from the tack test showed better mucoadhesive properties for the uncrosslinked precursor solutions used as controls (Table 3), compared to the crosslinked hydrogels. Confocal microscopy was not only useful for mucoadhesive studies but also provided insights into the stability of the hydrogels, which is a desired property of the formulation being developed. Together, these methods underscore the multifaceted nature of mucoadhesion and provide a framework for optimizing hydrogel formulations for practical applications.

### 3.9 Evaluation of the swelling properties of the crosslinked hydrogels

Traits of hydrogel-based drug delivery systems such as mechanical properties, diffusibility, capacity to retain and release the active pharmaceutical ingredients are influenced by their swelling properties. [56] We examined the swelling properties of the hydrogels at two different concentrations as shown in Figure 8. The hydrogels demonstrated concentration dependent swelling capacity with 2% HATA having the highest swelling ratio. Equilibrium swelling was reached after 24 hours post immersion in the PBS with pH of 7.4 imbibing about 61% of its original weight. On the contrary, 1% HATA hydrogels swelled minimally to about 6% of its original weight and decreased rapidly to 0.8% after the end of the experiment. These hydrogels with limited swelling capacity are categorized as low-swellable or non-swellable hydrogels [57] suitable for drug delivery applications. [58] The high swelling ratio is due to the increased polymer concentration which increases linearly with the number of hydrophilic chains available for water uptake. Also, the 1% HATA polymer’s ability to swell was influenced by the degree of crosslinking which might have been intense considering the low number of photocrosslinkable groups available for crosslinking with the same intensity of light therefore “over crosslinking” the chains resulting in tighter networks in comparison to the higher concentration of the polymer crosslinked with the same intensity of light.

**Figure 8.**
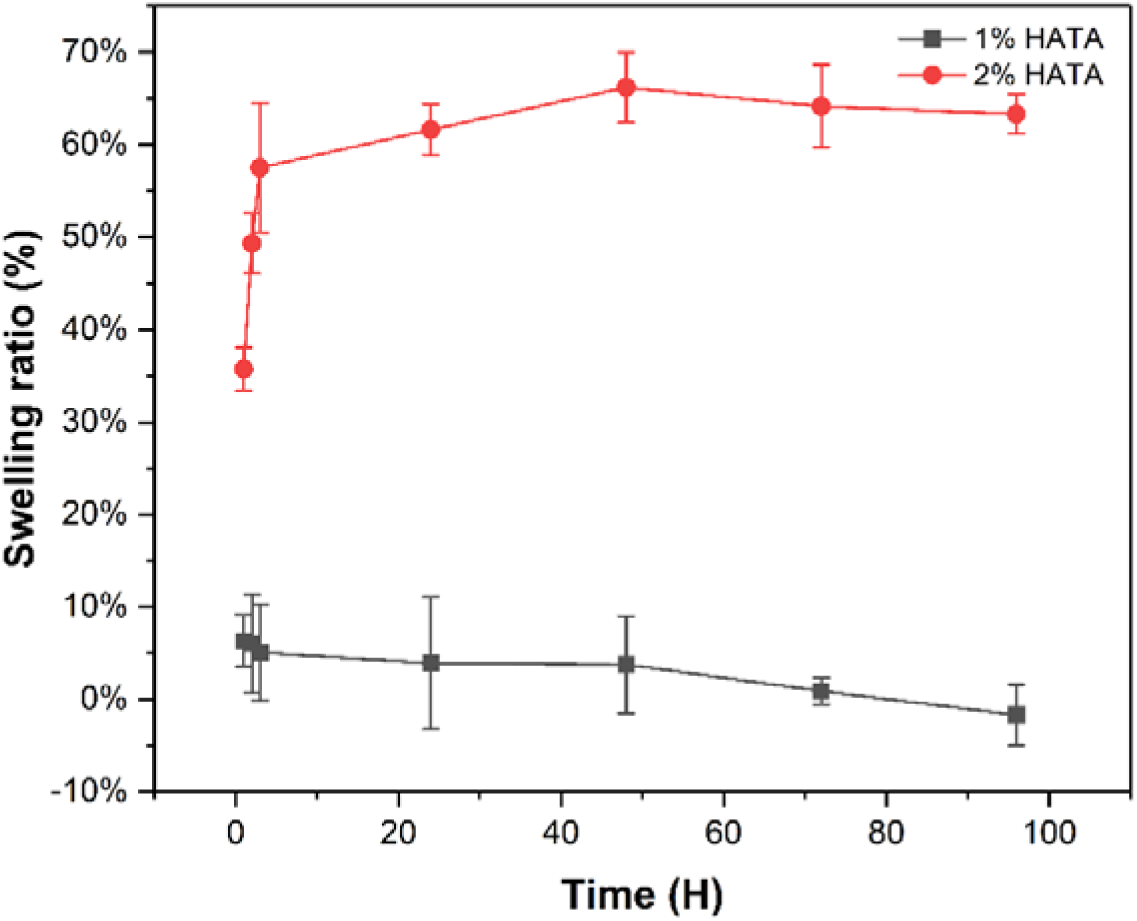
Swelling ratio of HATA hydrogels

**Figure 9.**
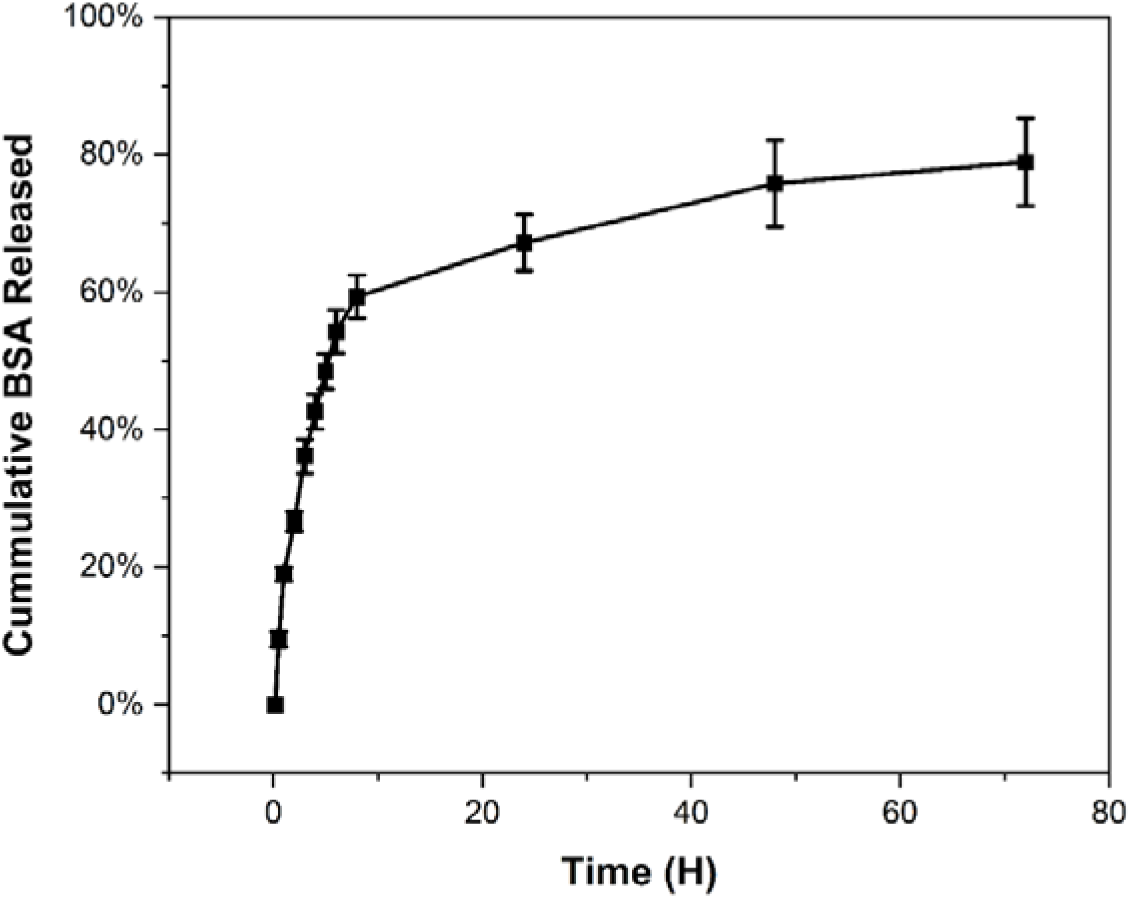
Cumulative release of BSA-rhodamine from 1 % w/v HATA hydrogel

### 3.10 BSA-Rhodamine release from hydrogel

Having established the criteria for the selection of suitable mucoadhesive DDS, we chose 1% HATA as a suitable candidate to be used as a DDS. Also, this formulation can be categorized as low swelling or non-swelling hydrogels making it a viable candidate among the other concentrations tested for drug delivery applications. The hydrogel demonstrated a rapid initial burst, releasing approximately 76% of the BSA within the first 24 hours as shown in Figure 10. Subsequently, the release rate plateaued remaining constant until a further 80 hours. The release curve was fitted with the Korsmeyer-Peppas model [59] to establish the mechanism underlying BSA-rhodamine release from the hydrogel. From the model, the k and the n values representing the kinetic constant and the release exponent were estimated as 0.3442 and 0.2048 respectively with coefficient of correlation (R^2^ = 0.939). With n < 0.5, BSA release from the hydrogel is purely diffusion-controlled and fulfils the Fickian diffusion criteria with minimal influence from matrix swelling or erosion as demonstrated in the significantly low swelling nature of the hydrogels. Compared to works already reported in the literature using dextran-based [60] or hyaluronic acid-based [61] hydrogels for BSA or a small molecule drug respectively, we observed a relatively high kinetic constant *(k)* further underpinning the rapid release of BSA in the early stage of the release studies. With hydrophilic systems such as HATA based hydrogels with no electrostatic or covalent interactions between the model protein and the polymer, it was not surprising to have observed the burst release of the model protein rapidly in the first 24 hours. Also, previous studies have estimated a significantly higher mesh size of HA-based hydrogels [57,62] which serves as a rapid escape route for BSA with diameter of 4.2 nm and 14.4 nm [63] in length which further explains the rapid release of the model protein from the hydrogels.

## 4 Conclusion

The present study reports the panoramic physicochemical characterization of photocrosslinked HATA-based hydrogels from precursor solutions to crosslinked hydrogels, prospectively applicable for nose-to-brain drug delivery via a medical device. Mucoadhesion was characterized extensively from the mechanical point of view using various physicochemical methods. By employing advanced confocal microscopy imaging, we emphasized the importance of crosslinking the hydrogels *in situ* by introducing photon count per pixel as a novel parameter to describe the stability and, most importantly, the mucoadhesive properties of the hydrogels on *ex vivo* porcine mucosal tissues. Unlike the classical wash-off methods which estimate indirectly the adhered materials on the mucosa surface, our novel method developed and designated parameter allowed direct quantification of the adhered hydrogels via advanced *in situ* microscopy imaging. We also highlighted the importance of the *in situ* crosslinking approach compared to classical application of uncrosslinked formulations, which are easily raided by mucociliary clearance. Additionally, nanoindentation was used as a novel technique to evaluate the local interfacial viscoelastic properties or effective Young’s modulus of the hydrogels, establishing a correlation between polymer concentration, irradiation duration, and their influence on mucoadhesive properties. This relation served as a yardstick in optimizing irradiation time and viscoelastic properties without compromising mucoadhesive properties of the formulation. Having thoroughly examined these physicochemical parameters, we then used BSA-conjugated rhodamine as a model protein drug and studied its release kinetics. Future experiments would seek to combine our drug delivery system with the nasal device already patented (EP 3865173 A1) for *in vivo* animal studies. We also aim to improve the release kinetics of hydrogels through the production of controlled release systems for protein therapeutics delivery.

## CRediT authorship contribution statement

**Seth Asamoah:** Writing – review & editing, Writing – original draft, Validation, Methodology, Investigation, Conceptualization. **Martin Pravda:** Supervision, Writing – review & editing, Methodology, Conceptualization. **Eva Šnejdrová:** Funding acquisition, Supervision, Writing – review & editing, Methodology, Conceptualization. **Martin** Č**epa:** Writing and Methodology. **Mrázek Ji**ř**í:** Writing and Methodology. **Carmen Gruber-Traub:** Funding acquisition, Supervision, Writing – review & editing. **Vladimír Velebný**: Supervision, Writing – review & editing

## Data availability

Data will be made available upon request.

