## Supplementary Material for "Photocrosslinked mucoadhesive hyaluronic acid hydrogel for transmucosal drug delivery"

Viscoelastic properties of hydrogels

Table S1: Linear viscoelastic region of the different concentrations of photocrosslinked HATA (Mean ± SD, n=3.)

| Polymer concentration (% w/v) | G' (Pa) | G'' (Pa) |
| --- | --- | --- |
| 1% HATA | 232.331 ± 9.935 | 0.869 ± 0.041 |
| 2% HATA | 348.851 ± 0.281 | 3.086 ± 0.065 |
| 3% HATA | 449.632 ± 1.097 | 14.600 ± 0.652 |

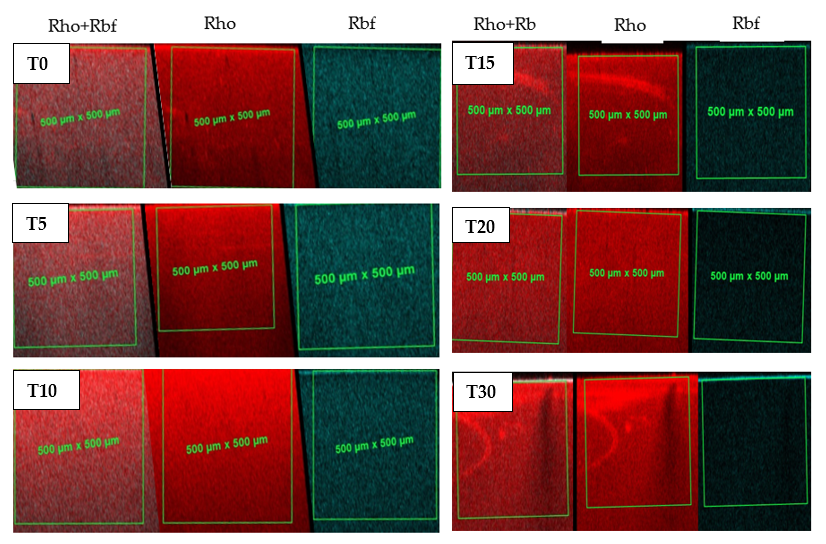

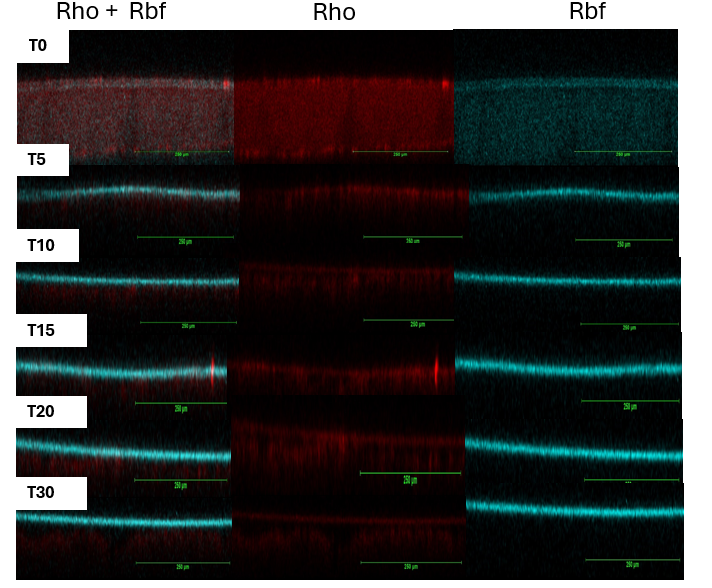

**A**

**B**

Figure 1S. A representative confocal microscopy image out of the triplicate measurements of rhodamine-HATA crosslinked hydrogel image **(A)** Uncrosslinked rhodamine-HATA image **(B).** Rho+Rbf: combined signal from riboflavin and rhodamine, Rho: rhodamine only signal and Rbf: riboflavin only signal. T0, T5, T10, T15, T20 and T30 represents time 0, 5, 10, 15, 20 and 30 minutes respectively.

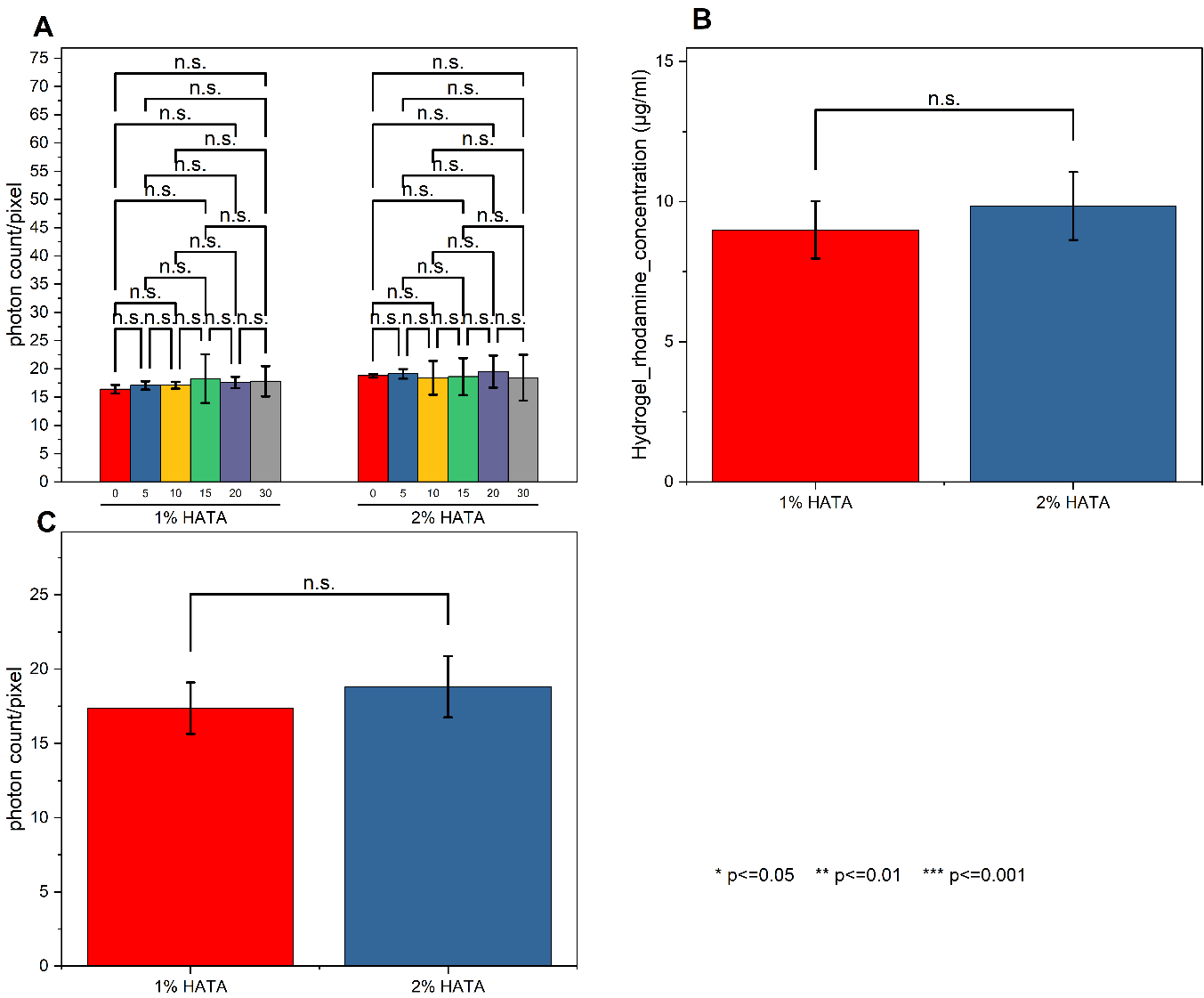

Figure 2S Crosslinked hydrogels photon count/pixel, comparison between different time points (i.e. 0, 5, 10, 15, 20 and 30 minutes post wash-off test) of the two different concentrations of 1% HATA and 2% HATA **(A)** Comparison of the two different concentration of the crosslinked hydrogels based on their rhodamine concentration (**B)** hydrogel photon count/pixel of the two different concentrations of polymer (**C)**
